# Cohort Profile: HealthWise Wales. A research register and data collection and analysis platform with linkage to NHS datasets in Wales

**DOI:** 10.1101/613547

**Authors:** Lisa Hurt, Pauline Ashfield-Watt, Julia Townson, Luke Heslop, Lauren Copeland, Mark Atkinson, Jeffrey Horton, Shantini Paranjothy

## Abstract

**Purpose:** Recruitment and follow-up in epidemiological studies is challenging, time-consuming and expensive. Combining online data collection with a register of individuals who agree to be contacted with information on research opportunities provides an efficient, cost-effective platform for population-based research. HealthWise Wales (HWW) aims to support researchers by recruiting a cohort of “research-ready” individuals; advertising relevant studies to these participants; providing access to cohort data for secondary analyses; and supporting data collection on specific topics that can be linked with healthcare data.

**Participants:** Adults (aged 16 and above) living or receiving their healthcare in Wales are eligible for inclusion. Participants consent to be followed-up every 6 months; for their details to be used to access their routinely-collected NHS records for research purposes; to be contacted about research projects in which they could participate; and to be informed about involvement or engagement opportunities. Data are collected using a web-based application, with new questionnaires added every six months. Data collection on socio-demographic and lifestyle factors is repeated at two-to-three year intervals. Recruitment is ongoing, with 21,779 active participants (alive and currently registered).

**Findings to date:** 99% of participants have complete information on age and sex, and 64% have completed questionnaires on socio-demographic and lifestyle factors. These data can be linked with national health databases within the Secure Anonymised Information Linkage (SAIL) databank, with 93% of participants matching a record in SAIL. HWW has facilitated recruitment of 43,826 participants to 15 different studies.

**Future plans:** The medium-term goal for the project is to enrol at least 50,000 adults. Recruitment strategies are being devised to achieve a study sample that closely models the population of Wales, with sufficient numbers in socio-demographic subgroups to allow for the selection of populations for research from those groups. Potential bio-sampling methods are also currently being explored.

## Introduction

High-income countries continue to face major public health challenges, including persistent inequalities in health and wellbeing and the complex needs of ageing populations (1, 2). Meeting these challenges requires a strong research infrastructure to ensure that high quality evidence is generated, for example, on preventing the onset and progression of non-communicable diseases and providing effective and efficient health and care services (3). Large-scale longitudinal studies are an essential resource for studying health and wellbeing throughout the life course. It is estimated that around 3.5% of the UK population are current or recent contributors to cohort studies (4). Using web-based technologies potentially makes recruitment and retention of subjects in such long-term studies less time-consuming and expensive (5). Combining online data collection with a register of individuals who have volunteered to be contacted with opportunities to take part in research also confers additional efficiency (such as SHARE Scotland (6)), and can create a platform to increase public involvement and engagement with research. Increasing awareness of the purpose of research and opportunities for participation should result in increased recruitment to research studies, better quality research to inform policy and practice, and ultimately improved population health outcomes (7).

Wales has a population of over three million people, within clearly defined geographical boundaries and with relatively low levels of migration in or out (8). It faces major challenges from a post-industrial legacy of socio-economic deprivation and a high prevalence of unhealthy behaviours (3, 9). High-quality, population-based research in this setting has already provided important evidence for policy and practice in the United Kingdom and beyond (10). HealthWise Wales (HWW) aims to provide an integrated cost-effective platform for conducting population-based research, by:

1. Establishing a cohort of “research-ready” individuals consented for re-contact;
2. Collecting longitudinal data from participants on self-reported exposures and outcomes; and
3. Using routinely-available healthcare data through record linkage (11, 12).

Overall, HWW plans to contribute to shape the health and wellbeing of future generations in Wales, and help the National Health Service (NHS) in Wales plan for the future.

## Cohort Description

### Setting

Recruitment into HWW is ongoing and dynamic, with individuals joining (or leaving) on a continuous basis and with varying levels of participation during their life course. Recruitment started during a pilot phase (March 2015 to February 2016), followed by a public launch on February 29^th^ 2016. Recruitment protocols have been designed to ensure representation across all areas of Wales. Overall, the distribution of HWW participants by residence is representative of Wales. For example, census data show that 67% of the Welsh population live in urban areas (defined as settlements of at least 10,000 people) (13), compared with 63% in HWW.

### Eligibility criteria and participant recruitment

Adults (aged 16 or above) who are usually resident or receive their healthcare in Wales are eligible to join, and are invited to be:

1. Followed up at regular intervals to obtain information about their health, wellbeing and specific exposures (such as behavioural risk factors), and allow record-linkage with their routinely-collected health records;
2. Entered onto a database of potential participants for research studies;
3. Contacted to take part in specific research studies;
4. Actively engaged and involved in dialogue to shape the priorities of the research programme.

Television, radio and social media advertising campaigns have been undertaken to issue an open invitation to potential participants to register. The project has been promoted at a wide range of events across Wales (for example, cultural events such as the Eisteddfod and agricultural shows such as the Royal Welsh and Anglesey shows) and in different settings (such as NHS hospitals, general practices, pharmacy outlets, and large employers). Mass postal mail-outs have also been piloted in one Health Board area, and there are plans to extend this method of communication about the project to other areas of Wales.

There are three core recruitment methods that are adapted for use as appropriate in different settings. Participants can give their consent to join the project through an online web application, which is accessed via the project’s website (www.healthwisewales.gov.wales, see Figure 1). They can also be recruited face-to-face using tablets or paper-based sign-up forms at events and various locations across Wales, or can give their consent to be contacted by individuals from the Participant Resource Centre at Cardiff University who can provide them with further information about HWW by email or telephone. Protocols describing the use of these recruitment methods and relevant study materials in various settings have been developed and have been implemented by HWW champions (members of the public who have volunteered to engage and involve other members of the public) and facilitators/research assistants (Health and Care Research Wales and NHS support and delivery staff). A range of recruitment and data collection strategies have also been developed for individuals who do not have internet access and/or may not have been exposed to the advertising campaigns. These have included study recruiters using mobile technologies with an internet connection to collect data at community-based locations, or telephone-based consent and data collection.

**Figure 1:**
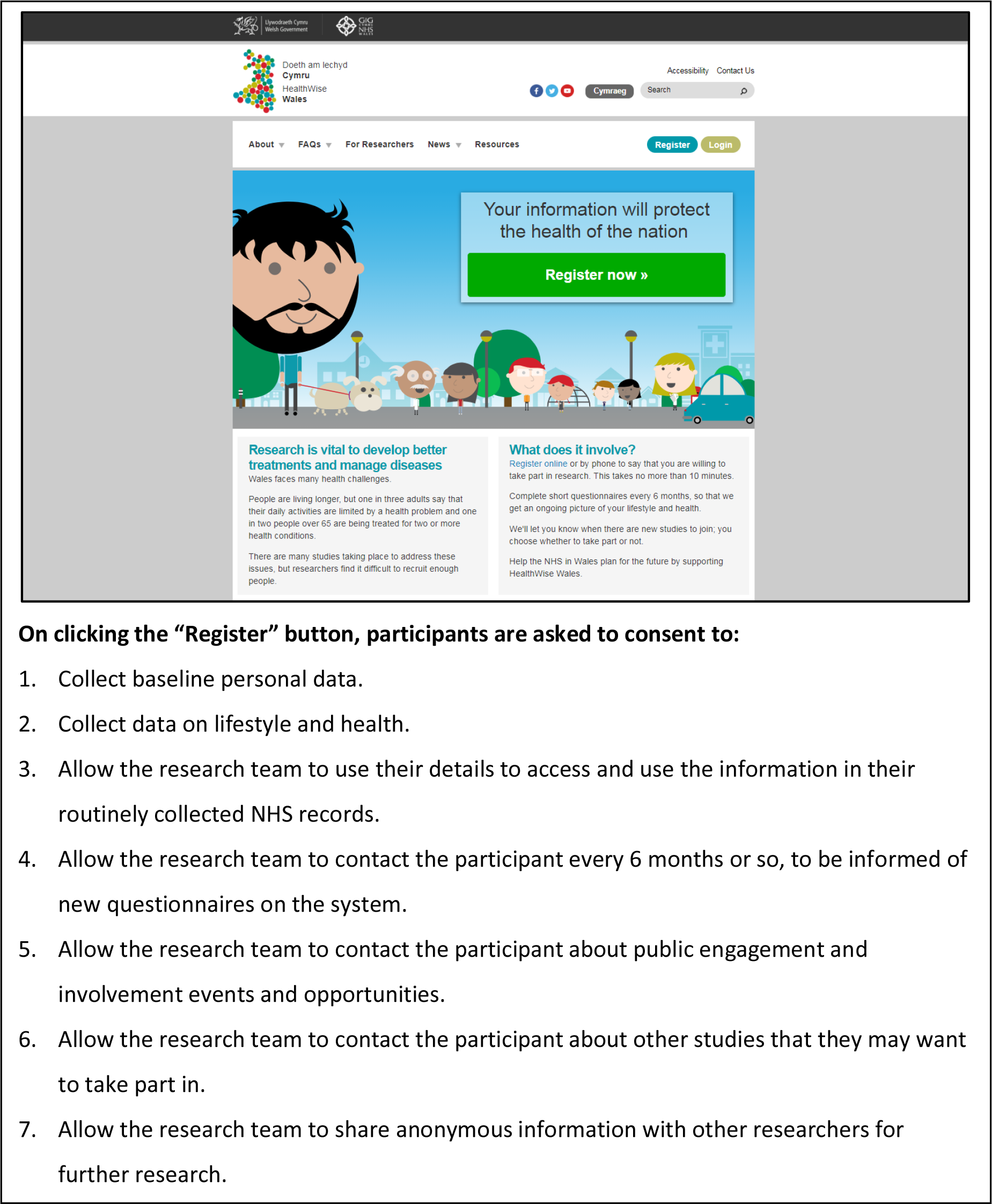
Website and Consent.

The medium term goal is to enrol at least 50,000 adults. This proposed sample size will be significantly larger than current population-based surveys in Wales, providing more precise estimates of the prevalence of exposures and outcomes in different socio-demographic groups, and adequate power to answer a range of different research questions about the determinants of health and wellbeing.

### Research themes

The project has five research themes:

1. Impact of social inequalities on health and wellbeing;
2. Environment, neighbourhood and health;
3. Maintenance of health and wellbeing in the working age population;
4. Wellbeing in later life; and
5. Innovation in health and social care services.

These themes are broad to guide data collection and facilitate use of the HWW platform by a wide range of health and social care researchers. Across these themes, there is a focus on four health areas (cancer, mental health, dementia and family life, pregnancy and early childhood health and development).

### Methods of data collection and follow-up

Data are collected using a web-based application, designed specifically for the project, which is accessible to participants through the main HWW website. New questionnaires are added every six months. These either collect information on items relevant to the research themes outlined above, or bespoke data to facilitate researcher-led projects that are aligned to the research themes. Descriptive information on the core research questionnaires, their availability to participants since the project launched in 2015, and completion numbers are presented in Table 1. These collect data on socio-demographic factors, lifestyle factors, home life, and mental health at baseline and will be repeated at two-to-three year intervals as appropriate. There is also an additional set of modified core questionnaires that collect information from pregnant women on their health and care.

**Table 1:**
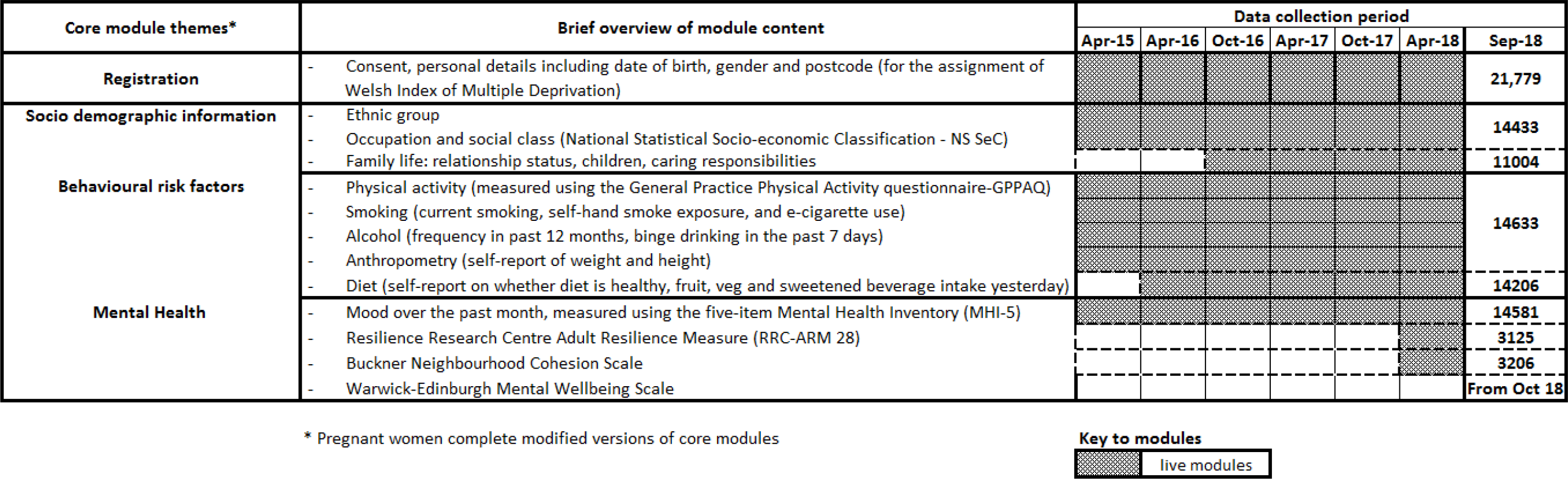
Outline of data collection questionnaires, timelines and summary of completions.

| Core module themes* | Brief overview of module content | Data collection period |  |  |  |  |  |  |
| --- | --- | --- | --- | --- | --- | --- | --- | --- |
|  |  | Apr-15 | Apr-16 | Oct-16 | Apr-17 | Oct-17 | Apr-18 | Sep-18 |
| **Registration** | - Consent, personal details including date of birth, gender and postcode (for the assignment of Welsh Index of Multiple Deprivation) |  |  |  |  |  |  | 21,779 |
| **Socio demographic information** | - Ethnic group |  |  |  |  |  |  | 14433 |
|  | - Occupation and social class (National Statistical Socio-economic Classification - NS SeC) |  |  |  |  |  |  |  |
|  | - Family life: relationship status, children, caring responsibilities |  |  |  |  |  |  | 11004 |
| **Behavioural risk factors** | - Physical activity (measured using the General Practice Physical Activity questionnaire-GPPAQ) |  |  |  |  |  |  | 14633 |
|  | - Smoking (current smoking, self-hand smoke exposure, and e-cigarette use) |  |  |  |  |  |  |  |
|  | - Alcohol (frequency in past 12 months, binge drinking in the past 7 days) |  |  |  |  |  |  |  |
|  | - Anthropometry (self-report of weight and height) |  |  |  |  |  |  |  |
|  | - Diet (self-report on whether diet is healthy, fruit, veg and sweetened beverage intake yesterday) |  |  |  |  |  |  | 14206 |
| **Mental Health** | - Mood over the past month, measured using the five-item Mental Health Inventory (MHI-5) |  |  |  |  |  |  | 14581 |
|  | - Resilience Research Centre Adult Resilience Measure (RRC-ARM 28) |  |  |  |  |  |  | 3125 |
|  | - Buckner Neighbourhood Cohesion Scale |  |  |  |  |  |  | 3206 |
|  | - Warwick-Edinburgh Mental Wellbeing Scale |  |  |  |  |  |  | From Oct 18 |

\* Pregnant women complete modified versions of core modules

**Key to modules**
|  |  |
| --- | --- |
|  | live modules |

Outcome data are obtained in two ways. First, data are collected on the HWW platform for patient-reported outcome measures and those relevant to conditions likely to be under-represented in routinely-collected data (for example, infections, metabolic diseases, psychiatric conditions and wellbeing). Second, outcome information can be obtained through record linkage with national health databases (such as the Patient Episode Database for Wales and general practice data) within the Secure Anonymised Information Linkage (SAIL) databank (11, 12). Future phases of the project will also include linkage with other administrative datasets.

The NHS Wales Informatics Service (NWIS, a trusted third party) uses the personal details of participants (with their consent) to generate an anonymised linking field (ALF_E) based on their name, address, gender and date of birth. This is used to link participants’ data with routinely-collected healthcare data sets, with 93% of active participants matching with a record in SAIL. The SAIL databank and Secure Access Portal and Protected HWW Information Repository (SAPPHIRe) are stored in separate areas of the UK Secure e-Research Platform (UKSeRP, (14)). Figure 2 shows the flow of project data, showing SAPPHIRe within UKSeRP where project-specific, anonymised HWW data can be accessed.

**Figure 2:**
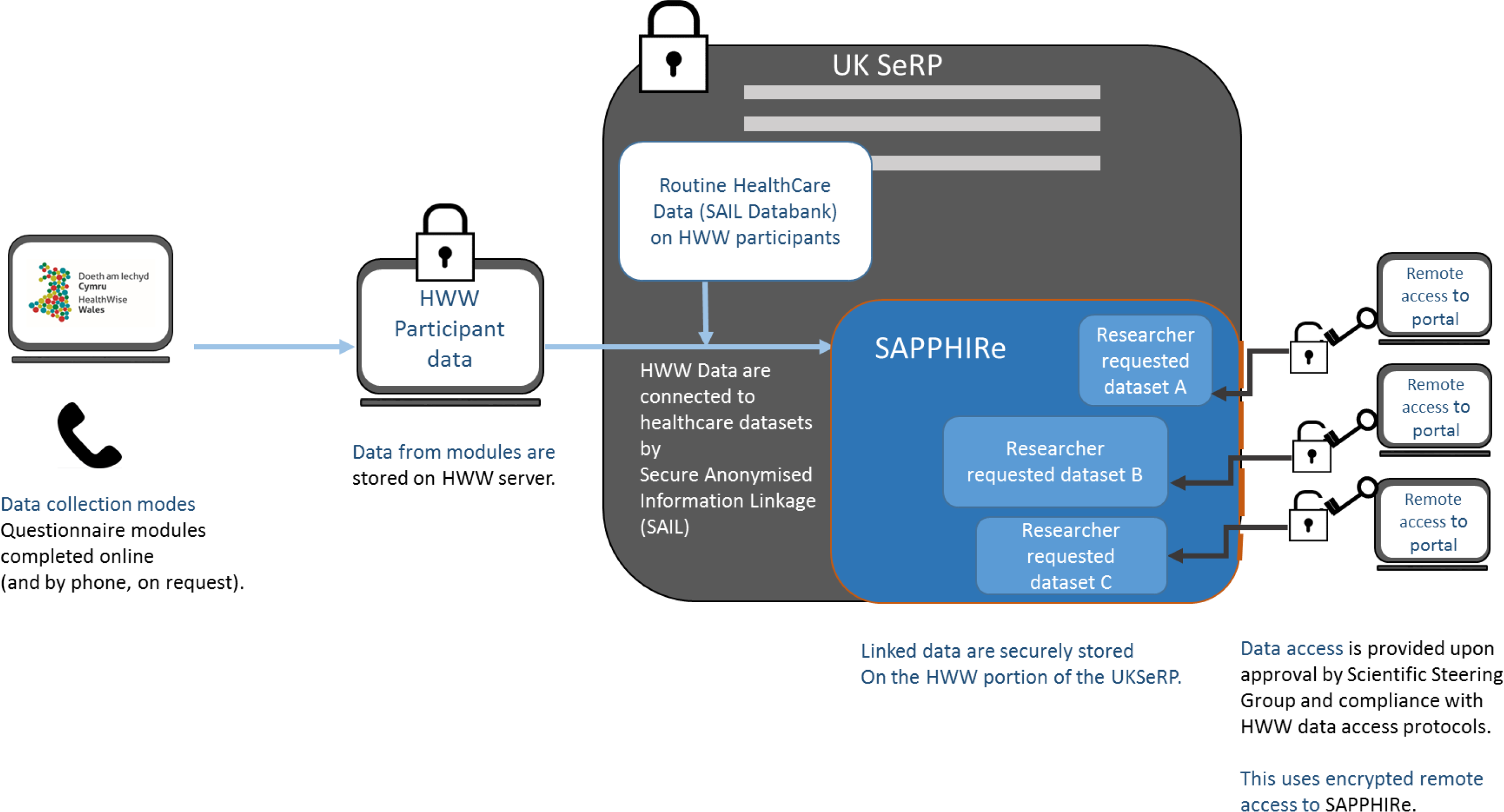
HealthWise Wales Data Flow.

### Characteristics of participants

There are currently more than 20,000 active participants (alive and currently registered). 99% of registered participants have complete information on age and sex, and at least 64% have completed the other core questionnaires. Table 2 shows the characteristics of active participants compared with data from published sources of Welsh data. Compared with the population of Wales, there is a higher percentage of participants who are 45 to 64 years old. The percentage of women is higher than in the general population (72% compared with 51%). The percentage of participants in non-white ethnic groups is the same as in the general population. 50 percent of participants are classified as being in higher managerial or professional occupations, compared with 27 percent of the population of Wales. In terms of health-related behaviours: 56 percent are classified as active or moderately active; 10 percent are current smokers (compared with 19 percent of the general population); and 50 percent drink more alcohol than recommended by UK guidance (compared with 40 percent of the general population). 28 percent of participants have a Mental Health Inventory score consistent with a common mental disorder and 32 percent have been diagnosed with or treated for a mental health condition (compared with 13 percent of the general population). Figure 3 shows the distribution of participants according to the Welsh Index of Multiple Deprivation compared with the population of Wales. There is a good representation of participants in each deprivation quintile, although a higher percentage of participants are from the least deprived quintile.

**Table 2:**
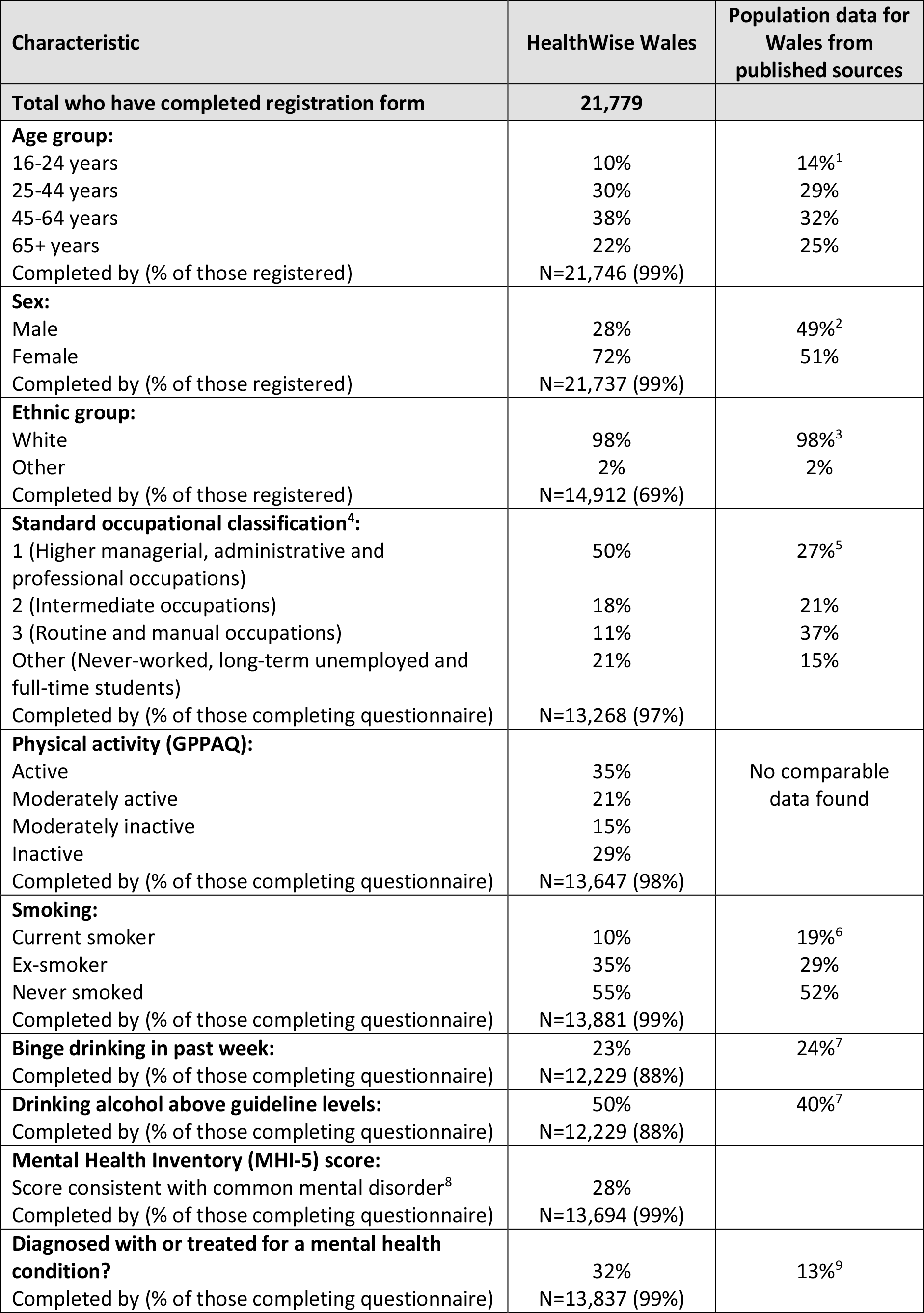
Characteristics of the HealthWise Wales cohort and population data from published sources for Wales.

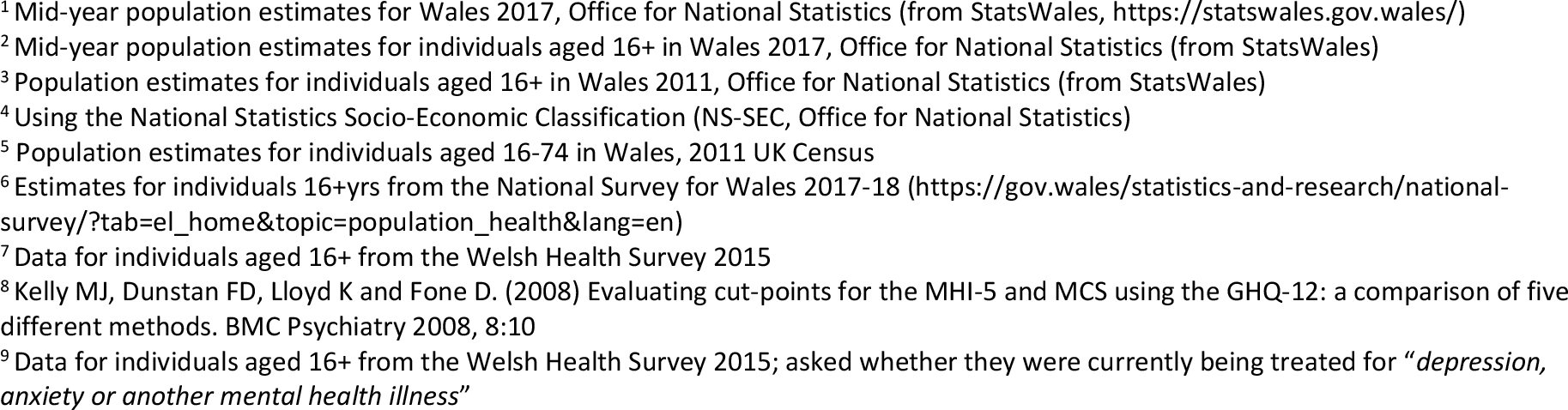

**Figure 3:**
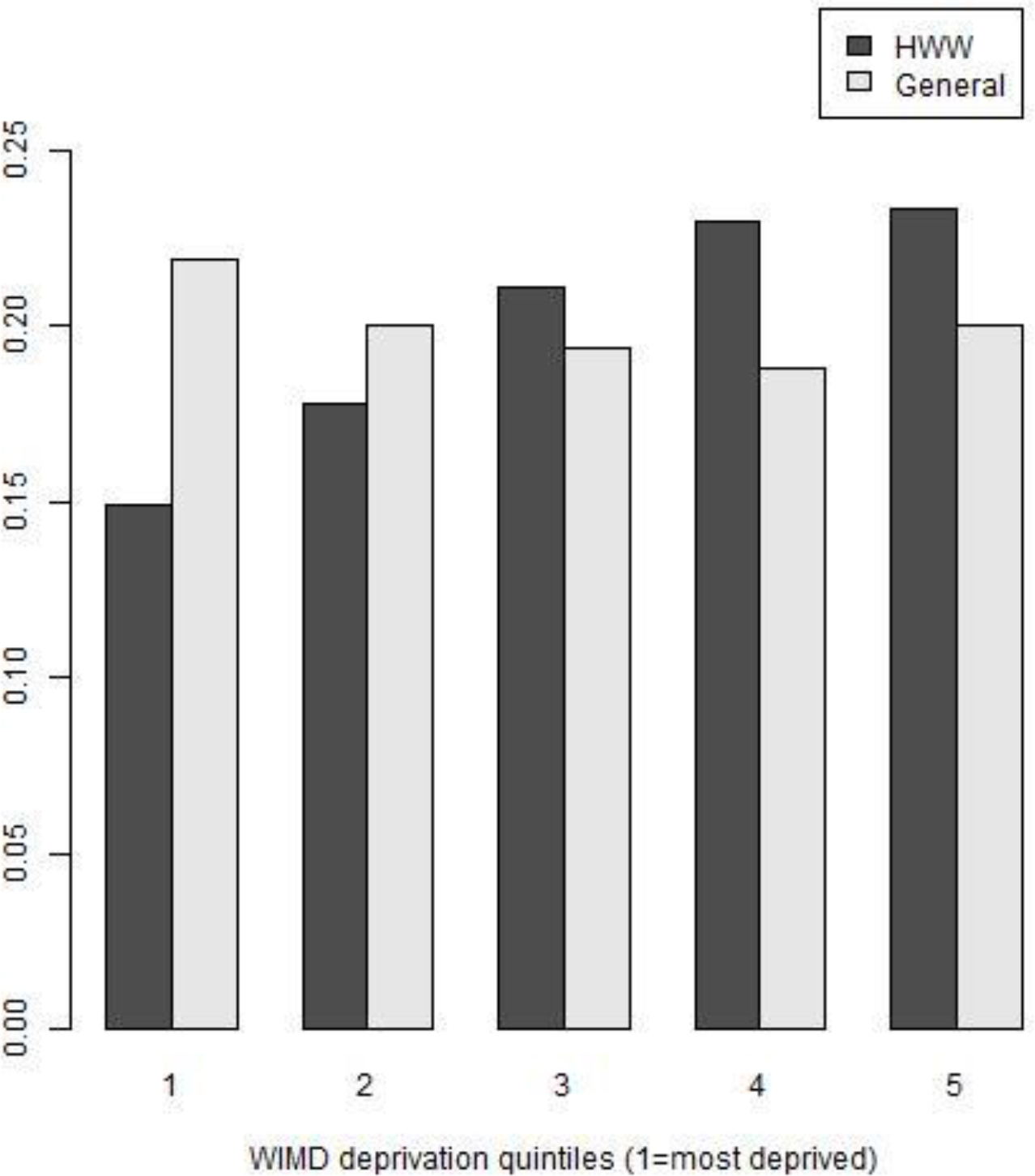
Proportion of participants resident in each quintile of the Welsh Index of Multiple Deprivation, compared with the general Welsh population.

### Patient and participant involvement

HWW has a specific focus on increasing public involvement and engagement in health and social care research. To ensure these aims are achieved, the project is overseen by a Public Involvement Delivery Board (PIDB), which is chaired by an independent member of the public and whose membership is predominantly comprised of members of the public. The PIDB provides scrutiny and assurance that the project is operating in the public interest, and provides advice and support in delivering best practice in accordance with the National Standards for Public Involvement (https://www.invo.org.uk/posttypepublication/national-standards-for-public-involvement/). The Board and the research team co-produced the project’s Patient and Public Involvement (PPI) policy and implementation plan. The research team has a dedicated PPI lead who is responsible for maintaining the policy document and ensuring compliance with it. All research team members are trained on facilitating public involvement. There are two PPI members of the research team, who have agreed objectives for their role and attend monthly meetings where they are actively involved in discussions and decision-making relating to research team activities. We have also trained 78 local health board members as facilitators to engage the public and recruit participants to HWW.

Involvement opportunities (including participation in media promotions or development and user-testing of data collection questionnaires) are regularly offered to participants through a quarterly e-newsletter. As a result, three participants became the faces of the advertising campaign in March 2017, others have participated in social media promotions, and 156 agreed to be members of a user-testing panel.

PPI is an essential criterion for all studies that use HWW, and researchers are required to describe the PPI they have undertaken when applying to use the data or the platform. PPI research team members scrutinise this element of applications as part of their overall assessment of all new projects.

### Ethical approval and governance arrangements

The project is overseen by an Executive Group, which provides oversight and decision making on the overall delivery of initiative, and receives advice from a Scientific Steering Group (SSG) and the Public Involvement Delivery Board. The role of the PIDB has been described above.

HWW received ethical approval from Wales Research Ethics Committee (REC) 3 on 16th March 2015 (reference 15/WA/0076). Substantial amendments are submitted when new questionnaires are added or if there is a substantial change to the content of participant-facing materials or recruitment model, in line with current guidance from the committee. The data collection system and study processes are designed to safeguard the integrity and confidentiality of data collected and generated for HWW research, and appropriate systems have been established and tested to report any failures in these respects. Standard Operating Procedures (SOPs) are in place to ensure that HWW is conducted within research governance regulations and compliant with the General Data Protection Regulation (GDPR) (EU) 2016/679. The research team meet with the HWW Data Guardian every six months to review the data governance processes in place and any matters arising.

### Funding

HWW is funded by Health and Care Research Wales.

## Research activities

HealthWise Wales supports researchers in three ways by: advertising relevant studies to participants; providing access to cohort data for secondary analyses via the researcher portal; and supporting data collection on specific topics within the platform that can then be linked with healthcare data. To date, seven studies have used the database to inform potential participants of an opportunity to take part in their research (see Table 3), with recruitment for each of these exceeding the required target. Nine studies have used the platform to collect data on study-specific questionnaires (see Table 4), with more than 5,000 participants providing data for each of these. In total, HWW has facilitated the recruitment of 43,826 participants to 15 different studies to date.

**Table 3:**
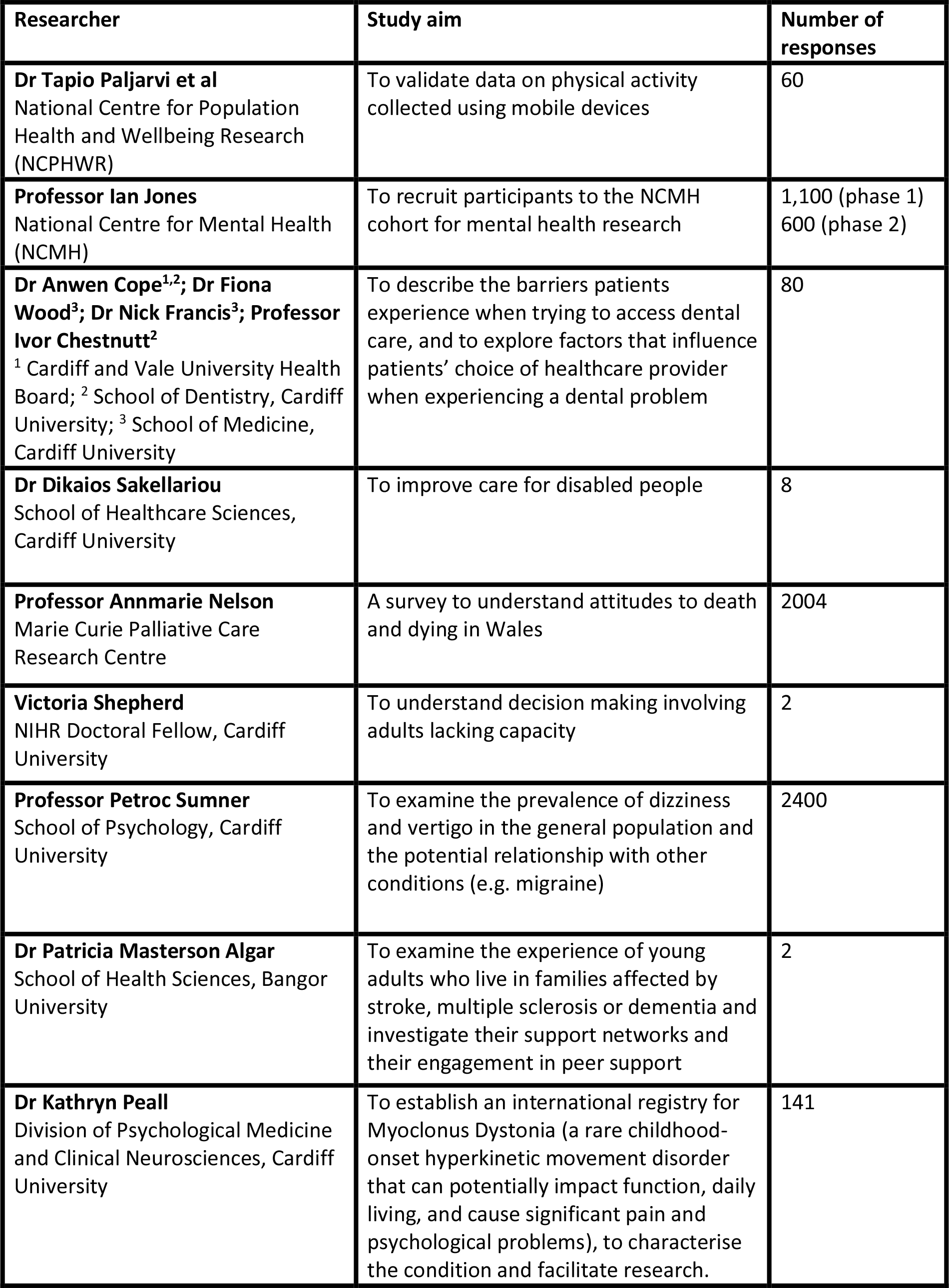
Engagement of HWW participants with research advertised via the HWW platform.

**Table 4:**
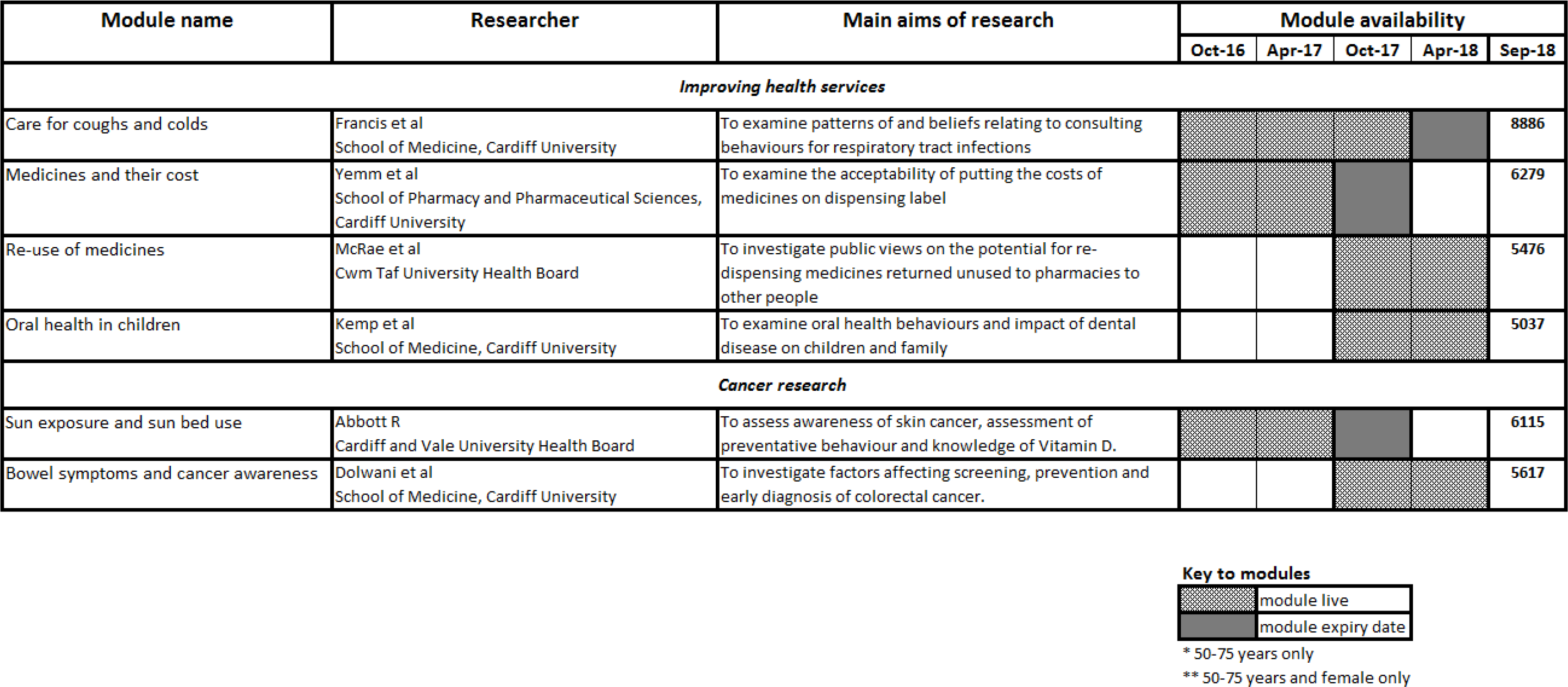
Examples of researcher-led questionnaire modules on the HealthWise Wales platform.

| Module name | Researcher | Main aims of research | Module availability |  |  |  |  |
| --- | --- | --- | --- | --- | --- | --- | --- |
|  |  |  | Oct-16 | Apr-17 | Oct-17 | Apr-18 | Sep-18 |
| Improving health services |  |  |  |  |  |  |  |
| Care for coughs and colds | Francis et al<br>School of Medicine, Cardiff University | To examine patterns of and beliefs relating to consulting behaviours for respiratory tract infections |  |  |  |  | 8886 |
| Medicines and their cost | Yemm et al<br>School of Pharmacy and Pharmaceutical Sciences, Cardiff University | To examine the acceptability of putting the costs of medicines on dispensing label |  |  |  |  | 6279 |
| Re-use of medicines | McRae et al<br>Cwm Taf University Health Board | To investigate public views on the potential for re-dispensing medicines returned unused to pharmacies to other people |  |  |  |  | 5476 |
| Oral health in children | Kemp et al<br>School of Medicine, Cardiff University | To examine oral health behaviours and impact of dental disease on children and family |  |  |  |  | 5037 |
| Cancer research |  |  |  |  |  |  |  |
| Sun exposure and sun bed use | Abbott R<br>Cardiff and Vale University Health Board | To assess awareness of skin cancer, assessment of preventative behaviour and knowledge of Vitamin D. |  |  |  |  | 6115 |
| Bowel symptoms and cancer awareness | Dolwani et al<br>School of Medicine, Cardiff University | To investigate factors affecting screening, prevention and early diagnosis of colorectal cancer. |  |  |  |  | 5617 |

**Key to modules**
|  |  |
| --- | --- |
|  | module live |
|  | module expiry date |

\* 50-75 years only
\*\* 50-75 years and female only

## Strengths and limitations

There are several strengths of HWW as a resource for research. More than 20,000 individuals with a diverse socio-demographic profile have already registered, and recruitment is ongoing. Matching rates of participant data with routinely-collected healthcare records are very high. In contrast with other population-based cohorts in the UK (4), HWW participants are younger, with most between 30 and 60 years old. This provides an opportunity to conduct longitudinal population studies with data collected pre-disease onset. Participants are also “research ready”; the examples given above demonstrate that the platform provides an effective way for the research community to reach an engaged, responsive cohort. A targeted retention plan is being developed with PPI representatives and a wider stakeholder group to encourage continued active participation in the project. Strategies found by other studies to be effective will be adapted to suit the HWW cohort, including the provision of real-time feedback to participants when they provide data, the development of an online community where participants can share their research experiences, and regular, diverse public engagement events to disseminate emerging results.

Men are currently under-represented in the cohort; only 28% of registered participants are male. Similarly, there are fewer individuals below 25 and over the age of 65 than in the general population, and a smaller percentage of participants from routine and manual occupations and in the most deprived wealth quintiles. Recruitment strategies to increase the number of participants in these groups are currently being devised. The aim is to achieve a study sample that closely models the population of Wales, with sufficient numbers in socio-demographic subgroups to allow for the selection of populations for research from those groups. For example, the cohort currently includes 5,000 men, providing a substantial sample size that will be adequate for some analyses.

Currently, bio-samples are not collected from participants. Formative research examining the willingness of individuals to provide different types of biological samples for research as part of their participation in HWW showed that 83% would be willing to do so. Options for a strategic approach to bio-sampling across Wales, and therefore a future enhancement that will increase the value of this cohort, are currently being explored.

## Collaboration

Figure 4 shows the application process for all research activities that can be undertaken using the HWW platform. All documentation informing researchers of how to apply to use the HWW platform was made public in June 2018, and access to the data has been possible since September 2018. A guide for researchers giving full details of the application and review process, and a copy of the application form, are available on the study website (www.healthwisewales.gov.wales/for-researchers).

**Figure 4:**
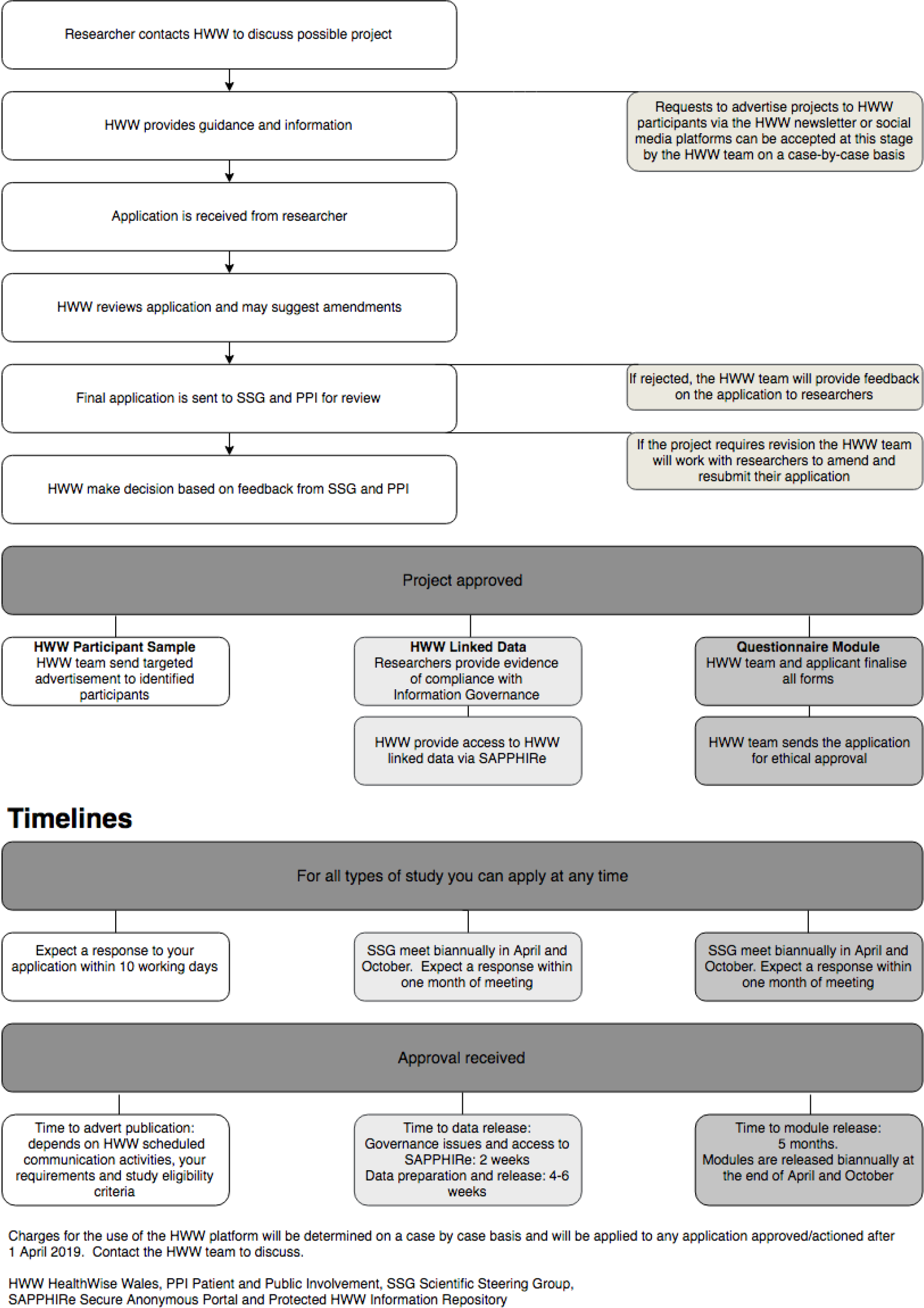
Flow diagram showing the application process for all HWW activities.

Requests to advertise projects to HWW participants via newsletters or social media are reviewed on a case-by-case basis by the HWW research team. The HWW ethical approval and participant consent permit HWW to advertise research projects to registered participants as long as they fit with the ethos and scope of the initiative. It is the responsibility of applicants to obtain ethical approval for the conduct of their specific study before HWW advertises it to participants. This ethical approval should specify that HWW will be used to help recruit participants.

Applications to use HWW for data collection or analysis are reviewed by the SSG and by PPI representatives, to assess that the project fits with the ethos of HWW, is scientifically sound, and that adequate PPI input has been sought in the development of the proposal. Once approved, researchers work closely with the HWW research team to deliver the project, including working together to prepare the application for a substantial amendment to the HWW ethical approval (which is needed for all new data collection). Researchers will need to provide evidence that they are bona fide researchers and have appropriate training in Research Data and Confidentiality procedures in order to gain access to the HWW data repository via SAPPHIRe.

Any publications related to the use of HWW resource must be sent to the research team, and a lay summary of the study findings must be shared with the team for publication on the website. All researchers using HWW should use the following standard text for acknowledgement in any publication arising from the use of the platform:

> *“This study was facilitated by HealthWise Wales, the Health and Care Research Wales initiative, which is led by Cardiff University in collaboration with SAIL, Swansea University.”*

Additional references are required for publications which use the SAIL databank and UKSeRP. These can be found in the Researcher Guidance document on the researcher tab of the HWW website: www.healthwisewales.gov.wales/for-researchers/

## Further details

Researchers should contact the research team (on or 0800 9172 172) before submitting their application to obtain guidance on how best to use the platform in their study, patient and public involvement processes, ethical requirements, questionnaire development, implementation and promotion.

## Conclusion

HWW is a research database of adults (aged 16 and above) living or receiving their healthcare in Wales that can support researchers by: advertising relevant studies to registered participants; providing access to cohort data for secondary analyses via the researcher portal; and supporting data collection on specific topics with record-linkage to healthcare data if required. It has been successful in recruiting a “research ready” cohort in Wales, and to date has facilitated recruitment of 43,826 participants into 15 studies.

## Acknowledgements

We gratefully acknowledge the contribution of Charlotte Bonner-Evans and Ameeta Richardson in the coordination, management and implementation of the platform. We would also like to thank Sean Dunn, Benjamin Dowie, Alex Coomber and others within the Participant Resource Centre (Cardiff University) and the Health and Care Research Wales Support and Delivery Centre for their contribution to recruitment and data collection, and the Welsh Government Communication team for their contribution to the Communications Plan. We acknowledge the substantial contribution of the Scientific Steering Group, the HealthWise Wales Executive group, the Public Involvement Delivery Board, Chris Stock, the Centre for Trials Research and Professor Mike Robling (Data Guardian). We also thank the participants.

